# Identification of novel genes regulating the development of the palate

**DOI:** 10.1101/2024.02.09.579685

**Authors:** Ashwin Bhaskar, Sophie Astrof

## Abstract

The International Mouse Phenotyping Consortium (IMPC) has generated thousands of knockout mouse lines, many of which exhibit embryonic or perinatal lethality. Using micro-computed tomography (micro-CT), the IMPC has created and publicly released 3D image datasets of embryos from these lethal and subviable lines. In this study, we leveraged this dataset to screen homozygous null mutants for anomalies in secondary palate development. We analyzed optical sections from 2,987 embryos at embryonic days E15.5 and E18.5, representing 484 homozygous mutant lines. Our analysis identified 45 novel genes implicated in palatogenesis. Gene set enrichment analysis highlighted biological processes and pathways relevant to palate development and uncovered 18 genes jointly regulating the development of the eye and the palate. These findings present a valuable resource for further research, offer novel insights into the molecular mechanisms underlying palatogenesis, and provide important context for understanding the etiology of rare human congenital disorders involving simultaneous malformations of the palate and other organs, including the eyes, ears, kidneys, and lungs.

## Introduction

Cleft palate with or without a cleft lip (known as orofacial clefts) is the most prevalent craniofacial birth defect, affecting approximately 1 in 800 babies annually (Rahimov et al., 2012). There is a significant difference in the incidence of orofacial clefting among different ethnic groups: the greatest incidence was found among Native Americans at 3.6 per 1000 births, followed by Japanese and Chinese populations, with a prevalence of 2.1 and 1.7 per 1000 births, respectively, while the lowest incidence is among African-Americans, at approximately 0.3 incidences per 1000 births (Wojcicki et al., 2016). Children born with this disorder have difficulties with feeding and speech when untreated. In developed countries, orofacial clefts are surgically treated early in childhood to close the clefts (Wojcicki et al., 2016). Despite the treatment, some affected individuals may still experience negative psychological, psychosocial, and financial hardships (Wehby and Cassell, 2010).

Palate formation proceeds in multiple phases (Hammond and Dixon, 2022; Som and Naidich, 2013). During the early stages of palate development, multi-potent neural crest (NC)-derived cells migrate to populate the first branchial arch, which subsequently gives rise to the maxillary and mandibular prominences. The secondary palate, the tissue that separates the nasal and oral cavities, forms when the paired outgrowths from the maxillary processes fuse at the midline and then with the primary palate, which is derived from the frontonasal prominence. In humans, palatal development takes place between weeks 4 and 10 of gestation (Mossey et al., 2009; Som and Naidich, 2013).

Due to the distinct developmental processes underlying the formation of the primary and secondary palate, several types of clefting are observed in patients and vertebrate model organisms: isolated cleft lip, isolated cleft palate, or clefting can be present both in the lip and palate (Mossey et al., 2009; Selleri and Rijli, 2023).

Mice are an established model for studying palate formation and clefting due to similarities in palate development between mice and humans (Selleri and Rijli, 2023). In mice, the growth of paired palatal shelves from the maxillary processes can be observed at embryonic day (E) 11.5. These processes undergo a dramatic increase in size through the proliferation of NC-derived mesenchyme and initially move downwards to be positioned on either side of the tongue by E13.5. The paired palatal shelves then elevate to become positioned horizontally above the tongue by E14.5. The fusion of palatal shelves at the midline begins soon afterward and results in the formation of a continuous shelf of the secondary palate. The fusion of the secondary palatal shelves is completed by E15.5, and the fusion of the primary and secondary palatal shelves is typically completed by E17.5 (Bush and Jiang, 2012).

The causes of cleft lip and palate in the human population are not completely understood. Epidemiological studies, genetic analyses, and modeling of lip and palate clefting in mice indicated that both environmental factors and genetic mutations are at play (Hammond and Dixon, 2022; Mossey et al., 2009; Selleri and Rijli, 2023). Among common mechanisms mediating palatal development in humans and mice are those regulated by Sonic Hedgehog (Shh) signaling (Cohen, 2010; Mansilla et al., 2006).

Modeling studies in mice indicated that precise levels of Shh gene expression in the NC-derived cells are essential for proper palate development, and both downregulation and upregulation of Shh in the NC-derived cells cause clefting and other defects in facial development (Jeong et al., 2004; Li et al., 2017). Additional players important for palate development include genes encoding secreted growth factors such as BMPs, Wnts, and Fgfs, and transcriptional regulators such as the Pbx family of transcription factors and *Snail1* (Ferretti et al., 2011; Losa et al., 2018; Welsh et al., 2018). However, much still needs to be learned. Therefore, it is critical to identify additional genes and their roles in palate development to broaden our understanding of mechanisms leading to the clefting of the lip and palate and to create more effective prognostic, diagnostic, and treatment options. To discover additional genes regulating the formation of the secondary palate, we examined images of mouse mutants generated and imaged by the International Mouse Phenotyping Consortium (IMPC) using micro-computed tomography (micro-CT) and looked for the presence of clefting in the secondary palate at E15.5 and E18.5 in coronal optical sections. Our analyses revealed novel genes regulating palatal development and provided new insights into the genetic etiology of rare human birth defects jointly affecting the development of the palate and other organs.

## Materials and Methods

### Image Analysis

To identify novel genes regulating the development of the secondary palate, we screened E15.5 and E18.5 homozygous-null mutant embryos generated and imaged by the IMPC: https://www.mousephenotype.org (Groza et al., 2023). Coronal optical sections of micro-CT images were analyzed using the tools embedded within the IMPC website to identify embryos with defective palates. Embryos at E18.5 were preferred for analysis due to the palate being fully fused at this stage in wild-type mouse embryos. E15.5 embryos were also analyzed. In E15.5 controls, the anterior-most and the posterior palate were often not yet fused. Therefore, cleft palate was only marked in those E15.5 mutants in which the palatal shelves were not fused throughout the entire length of the palate. Mutants containing cleft palates were also evaluated for eye defects, a category suggested by gene set enrichment analyses using Metascape (see Results and Discussion).

### Statistical Analysis

According to the Centers for Disease Control, the incidence of cleft palate in the human population is 1 in 1700 (Mai et al., 2019). Of 100 E18.5 wild-type embryos examined in this study, only one had a cleft palate (wild-type embryo ID: X63574-04). Using the binomial probability formula, we obtained a value of p=0.0554948102 (rounded up to 0.06 or 6%), an incidence of cleft palate expected solely by chance. Among the identified novel mutants with cleft palate, homozygous-null mutants in the *Spop* gene exhibited the smallest penetrance of cleft palate; only 1 of 11 *Spop*-null embryos had cleft palate. The binomial probability of observing 1 cleft palate out of 11 individuals is 0.007, indicating that the presence of a cleft palate was unlikely by chance alone, even at such low penetrance. The cutoff would be 1 cleft palate in 80 embryos (p=0.047).

### Identification of novel cleft palate genes

To find which of the cleft palate mutations were novel, we queried Mammalian Phenotype ontology terms associated with the genes on our list (Table 1). To do so, we queried the list of genes in Supplemental Table 1 against mammalian phenotype terms (https://www.informatics.jax.org/batch), Supplemental Tables 2A-C. In addition, we compared the genes on our list with those in a cleft gene database (https://bioinfo.uth.edu). Genes not previously identified as playing a role in palatal development were considered novel (Table 1, red ink). Mutants with cleft palates that were identified by the IMPC but otherwise not studied were marked in purple (Table 1).

**Table 1.**
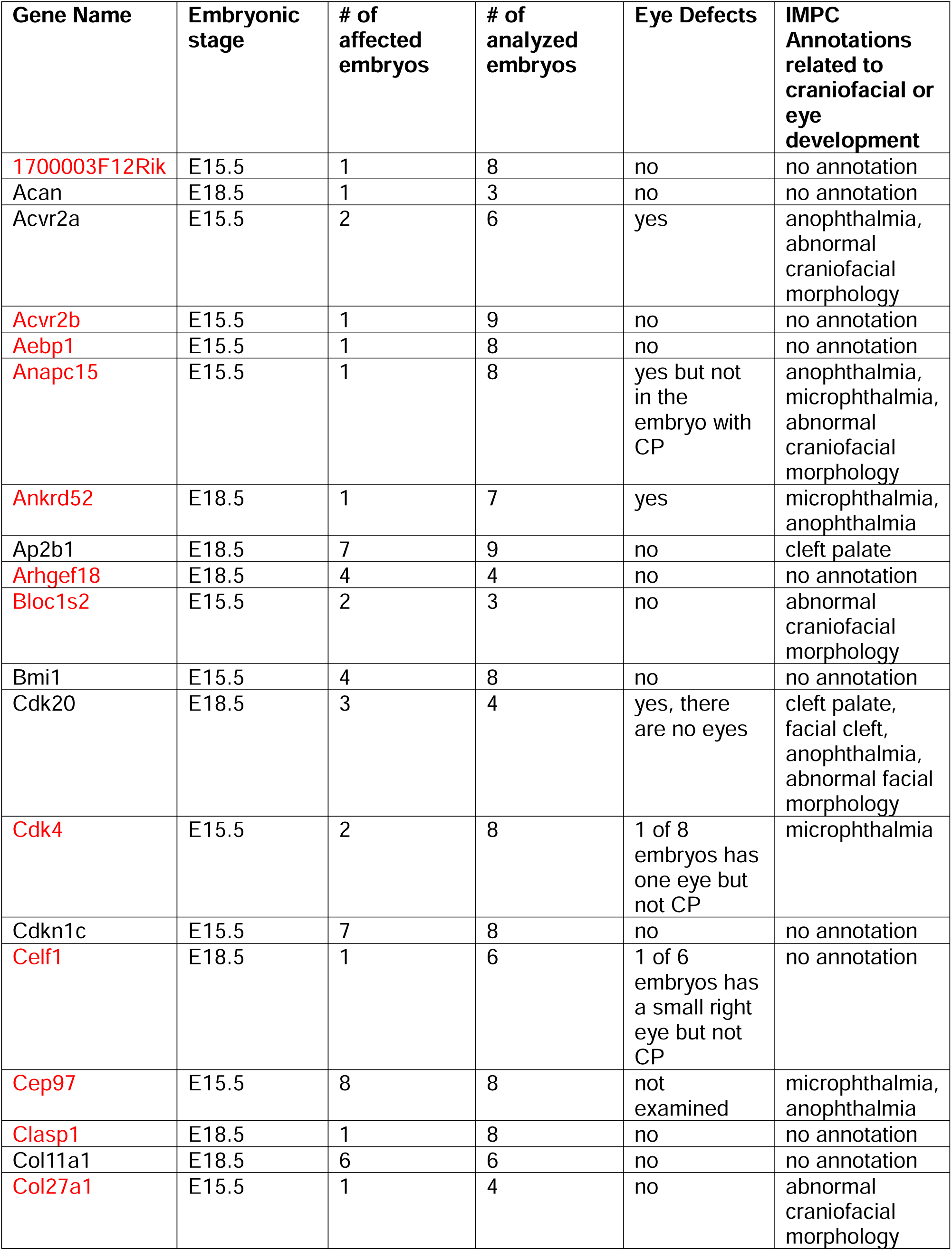
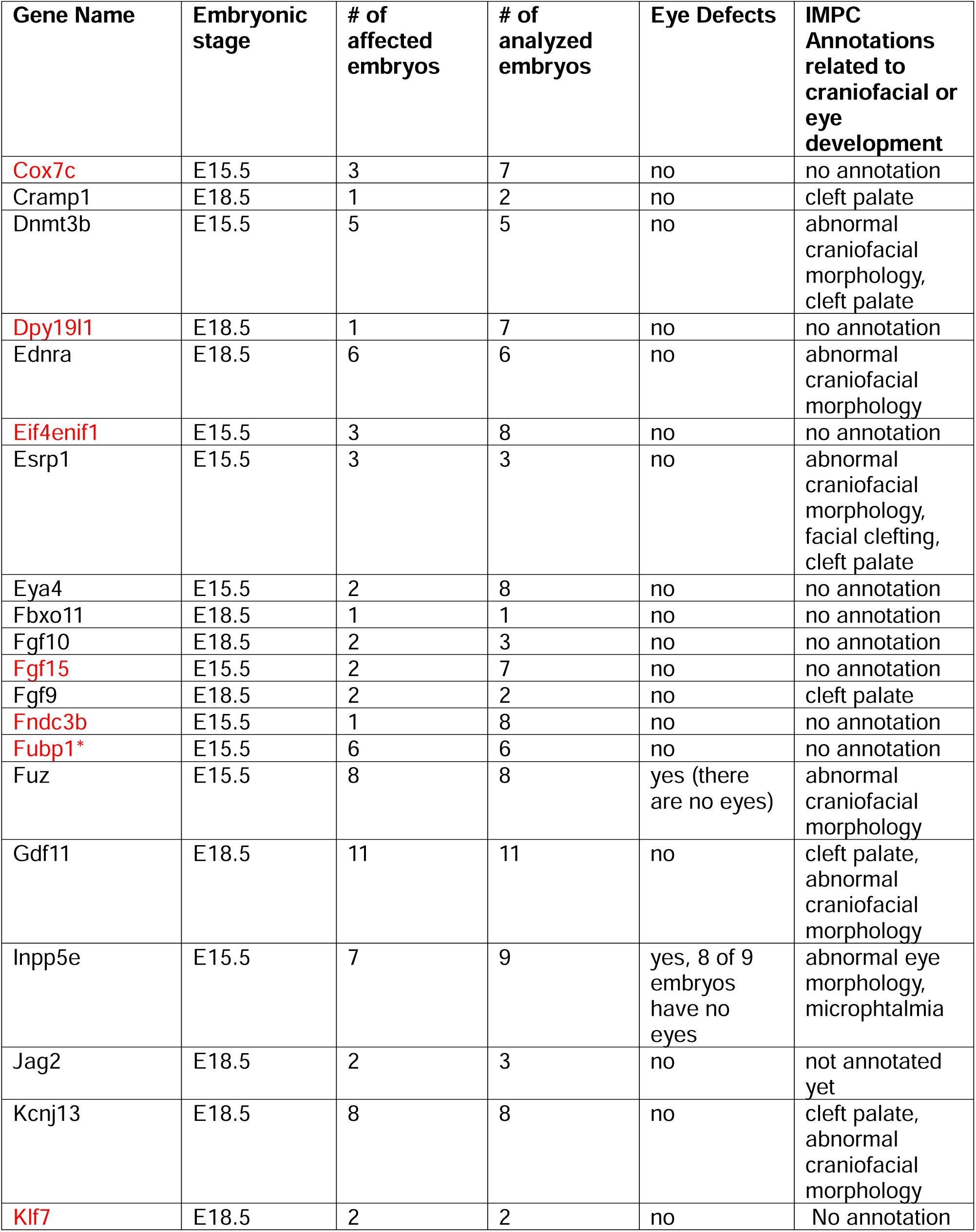
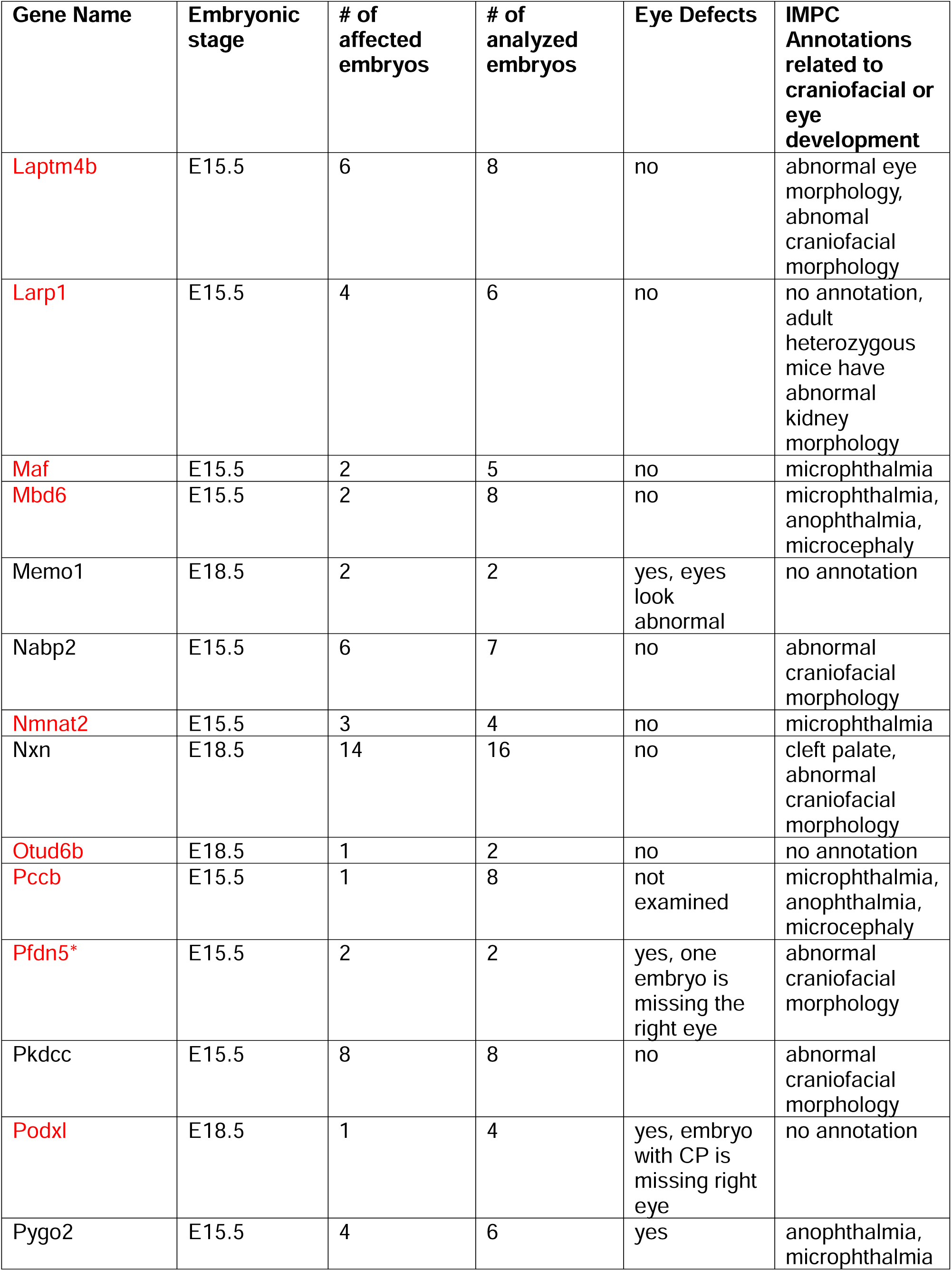
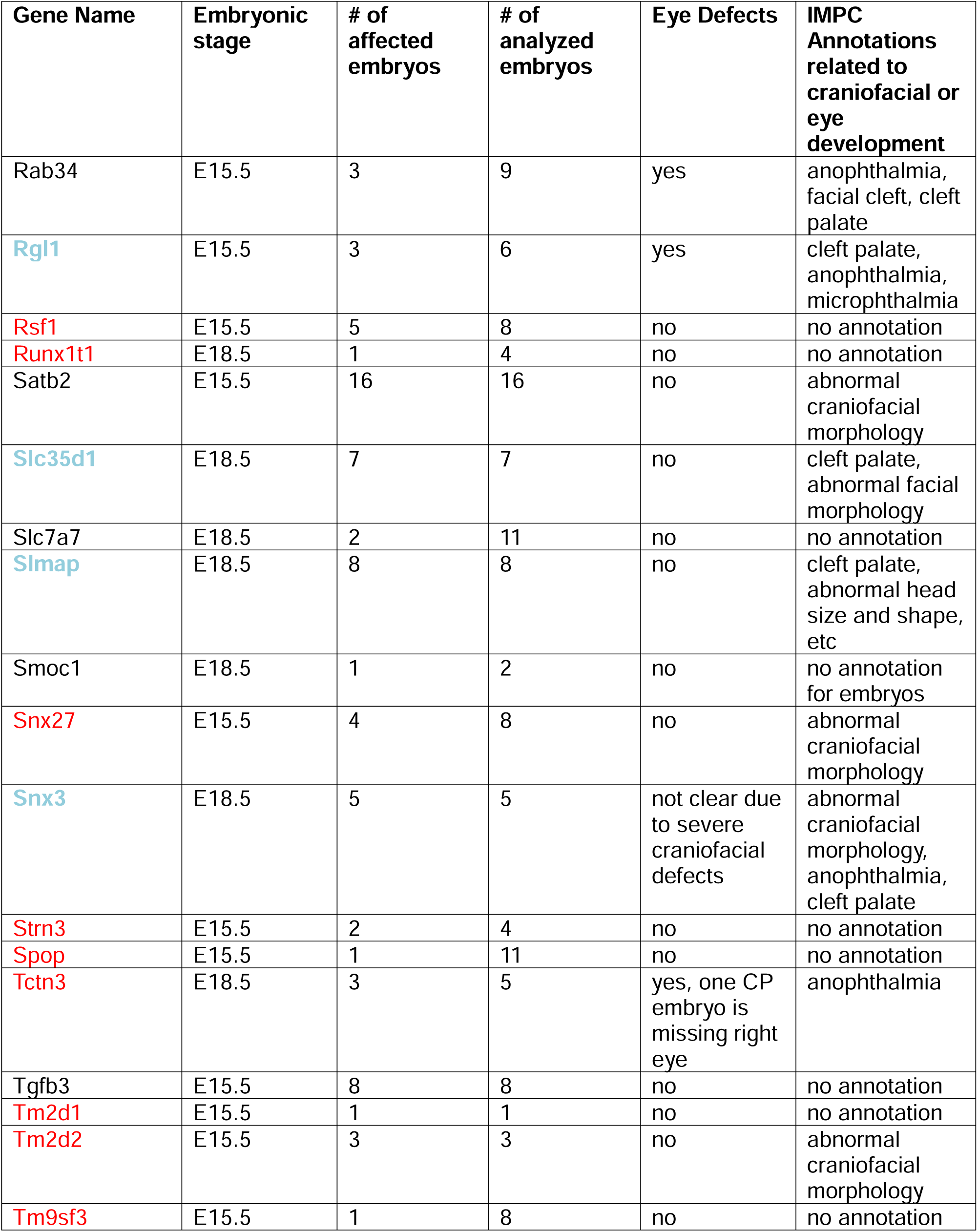
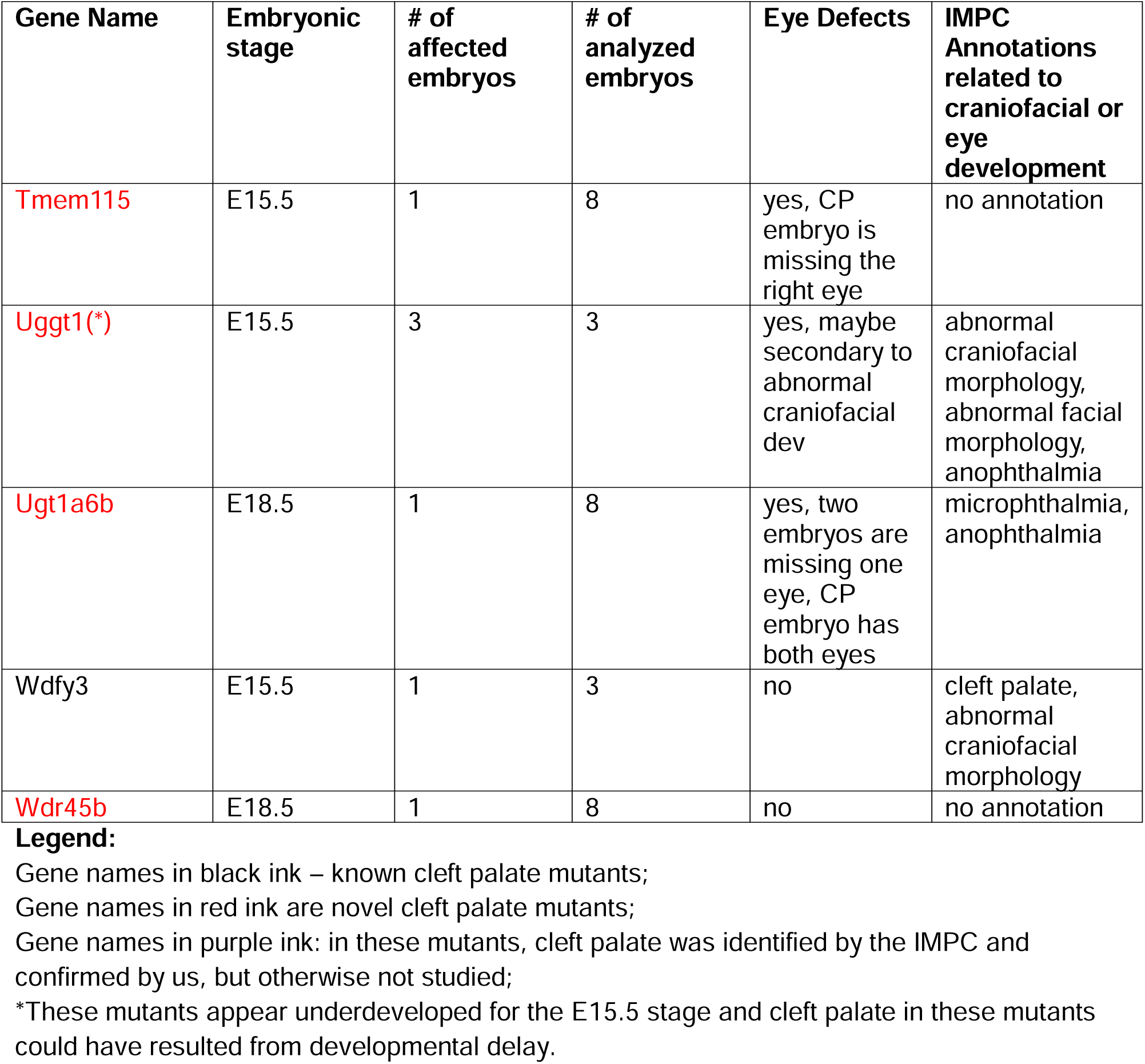
Incidence of cleft palate in the analyzed IMPC mutants.

### Pathway enrichment analyses

We used Metascape (Zhou et al., 2019) to identify enriched biological processes and pathways (Supplemental Table 3).

## Results and discussion

We examined the palatal morphology by analyzing coronal optical sections in 2987 embryos from 484 homozygous mutant lines generated and imaged by the IMPC (see Supplemental Table 1 for phenotype annotations of each examined embryo). Out of 484 homozygous mutant lines, 76 lines contained embryos with cleft palates (Table 1). To determine which mutant lines represented novel cleft palate genes, we used gene ontology terms and the Mouse Genome Informatics (MGI) along with the cleft palate database (https://bioinfo.uth.edu/CleftGeneDB/). These analyses indicated that of 76 genes, homozygous-null mutations in 45 were not previously associated with cleft palate (Table 1 and Supplemental Table 2). Four homozygous mutant lines in this set (*Rgl1, Slc35d1, Slmap*, and *Snx3*) were annotated as having cleft palates by the IMPC team, and we have also detected cleft palates in these mutants (Table 1, Figures 1-2). 7 of 484 mutant lines (*Cacna1s, Dhcr7, Fbxo30, Focad, Kdelr2, Robo1, Ryr1*) were previously associated with cleft palate in mice while the null mutants of these genes generated by the IMPC did not (Supplemental Table 3). Conversely, some of the mutants, even those displaying cleft palates at 100% penetrance (e.g. *Arhgef18,* a novel cleft palate gene) were either missed or not yet annotated by the IMPC as having cleft palates (Table 1). Therefore, although laborious, custom annotations of structural defects in the IMPC mutants are valuable.

**Figure 1.**
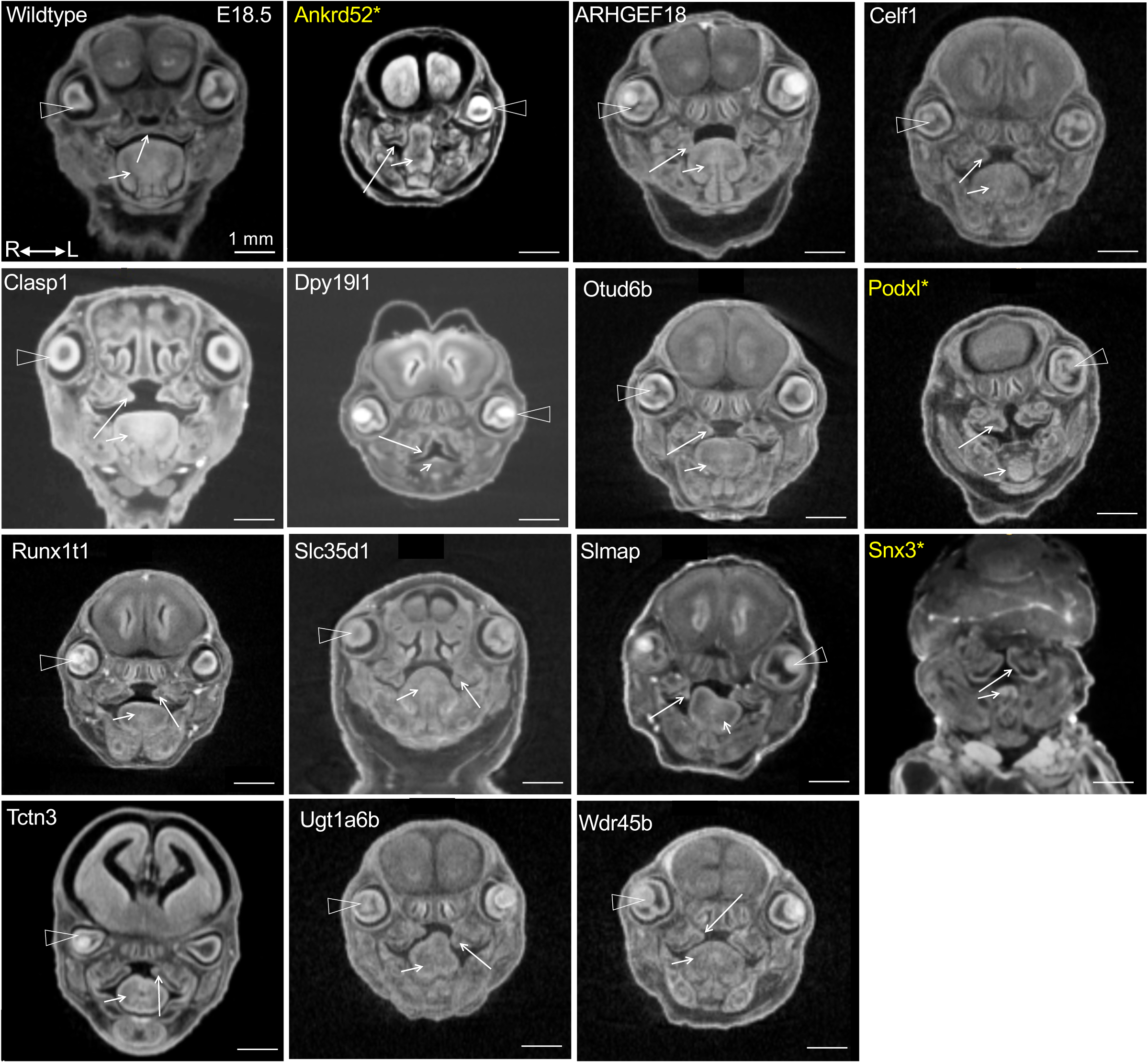
E18.5 mutants with cleft palate. Snapshots of coronal optical sections from micro-CT images at the eye level. Arrowheads mark eyes. Long arrows point at palatal shelves. Short arrows point at the tongue. Star and yellow font indicate mutants with palate and eye defects. Tables 1-2 contain additional information about the displayed embryos. All scale bars are 1 mm.

**Figure 2.**
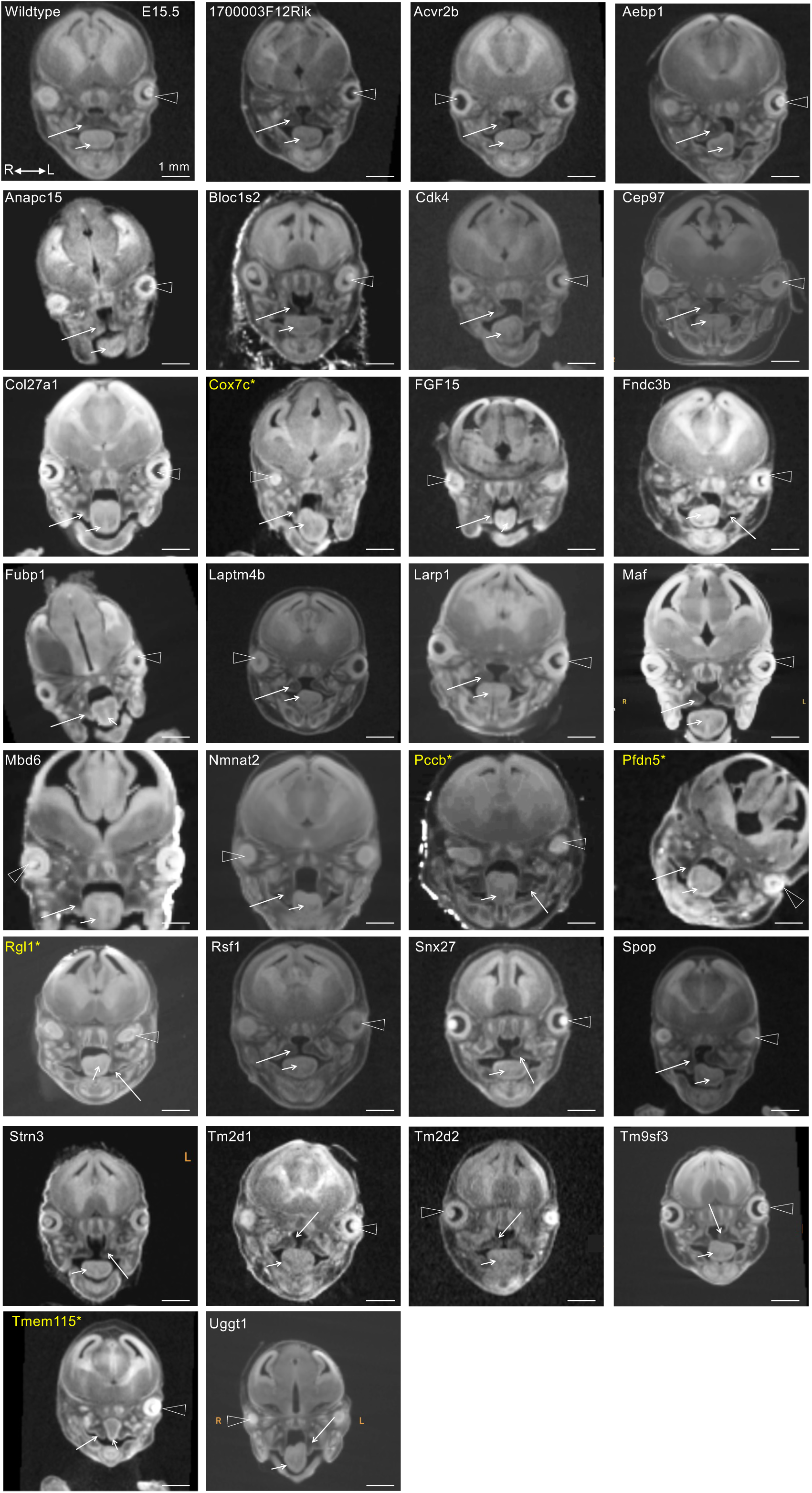
E15.5 mutants with cleft palate. Snapshots of coronal optical sections from micro-CT images were taken at eye level. Arrowheads marked eyes. Long arrows point at palatal shelves. Star and yellow font indicate mutants with palate and eye defects. Short arrows point at the tongue. Tables 1 and 3 contain additional information about the displayed embryos. All scale bars are 1 mm.

**Table 2:**
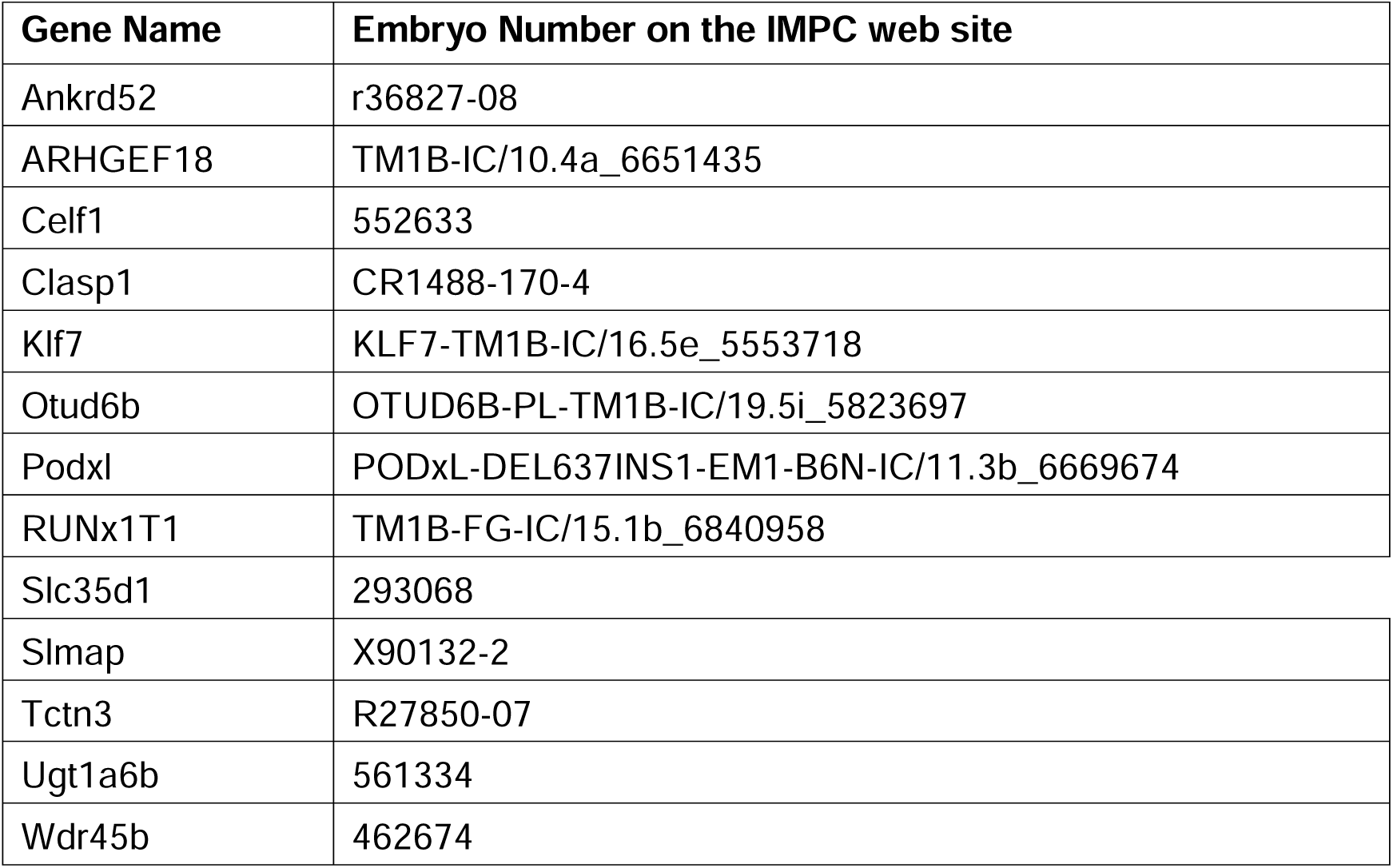
Embryos displayed in Figure 1 (E18.5)

**Table 3.**
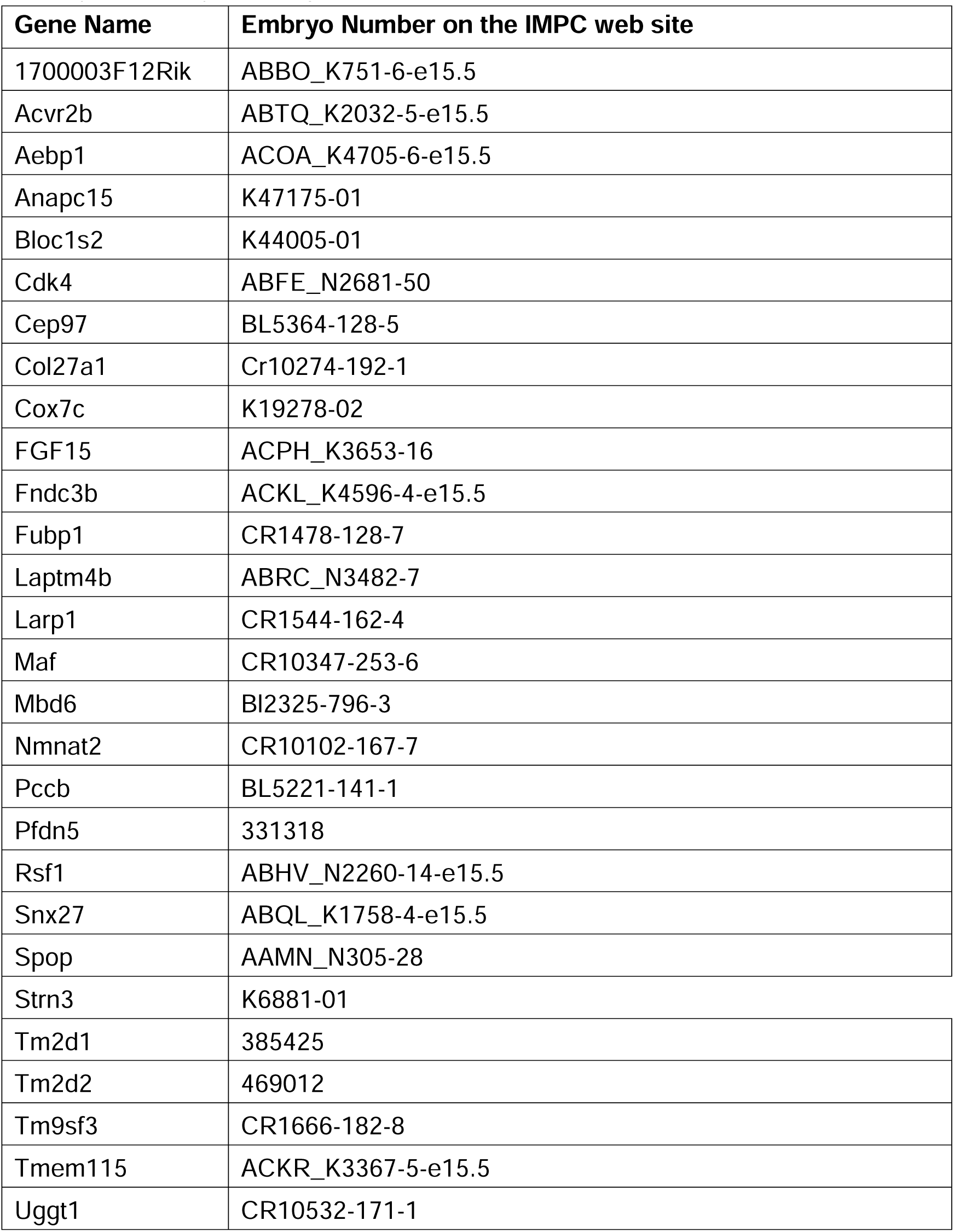
Embryos displayed in Figure 2 (E15.5)

Coronal sections displaying defective palatal morphology from 41 novel mutants annotated by us (red in Table 1) and 4 additional mutants annotated by the IMPC (purple, Table 1) are shown in Figures 1-2, with specific embryo information in Tables 2 and 3. 40 of the 41 novel genes were not previously associated with cleft palate phenotypes according to both the MGI and the CleftGeneDB databases. The *Col27a1* gene was associated with the cleft palate MP term in the MGI database but not in the CleftGeneDB database. Mutant *Col27a1* alleles were also generated independently of the IMPC; however, palate development was not examined in these mice (Plumb et al., 2011). And even though mutations in *Col27a1* were found in some patients with Steel syndrome whose symptoms included cleft palate (Girisha et al., 2022), no studies directly linked mutations in *Col27a1* to cleft palate. Therefore, we considered *Col27a1* a novel cleft palate gene. A gene that was initially considered a novel cleft palate gene but ultimately not considered novel in our analysis was *Bmi1*. *Bmi1* was not associated with cleft palate MP terms in the MGI database, but it was present in the CleftGeneDB. Upon further literature review, we found a statement, albeit without supporting image data, that 2 of 6 *Bmi1*-null embryos had cleft palate (Akasaka et al., 2001), therefore, we excluded *Bmi1* from the list of novel cleft palate genes.

Despite the uniform genetic background (C57BL/6N) used by the IMPC to generate null mutants, the presence of cleft palate was not fully penetrant in the majority of the homozygous null lines in which at least one case of cleft palate was observed (Table 1). Interestingly, even homozygous-null mutations previously known to be associated with cleft-palate phenotype in the literature displayed non-penetrant clefting phenotypes in the IMPC mutants (Table 1, e.g., *Bmi1*). Using a background incidence of cleft palate of 1 in 1700 (Mai et al., 2019), the presence of one case of cleft palate in 11 mutants per line was considered statistically significant (p=0.007). However, studying the mechanisms underlying palate clefting in mutants with low penetrance of palatal clefts is challenging. Although incomplete penetrance of structural defects in the IMPC mutants was observed previously, the underlying reasons for phenotypic variability in homozygous-null mutations on a uniform genetic background are complex and not fully understood (Dickinson et al., 2016).

We used the list of cleft palate-associated genes (Table 1) as input for pathway enrichment analysis via Metascape (Zhou et al., 2019), a tool that integrates ontology resources such as *KEGG Pathway, Reactome Gene Sets, GO Biological Processes, Canonical Pathways, CORUM*, and *WikiPathways*. This analysis identified enriched pathways regulated by cleft palate genes identified in our screen. Not surprisingly, “Skeletal system development“ was the top, most significantly enriched category (q= 4.20E-08, q is the probability corrected for multiple testing). Among the 20 genes in this category, *Acvr2b, Slc35d1*, and *Col27a1* were novel cleft palate genes identified in this study.

NC-derived cells give rise to the palatal mesenchyme, craniofacial bones, and cartilage; hence, we expected that pathways related to NC development would be enriched in our dataset. These pathways included: “Roof of mouth development” (q = 5.89E-07) and “sensory organ development” (q = 2.56E-05). Among the 31 genes in these two categories, *Acvr2b, Maf, Pfdn5, Fndc3b,* and *Col27a1* were novel (Supplemental Table 3).

Since NC-derived cells comprise the bulk of the palatal shelf tissue, it is reasonable to hypothesize that the development of organs regulated by the NC would also be affected in mutants with cleft palates. One such organ is the eye (Bryan et al., 2020; Gage et al., 2005; Williams and Bohnsack, 2015). Although ocular defects are among the top 30 birth defects that co-occur in patients with cleft palate (Sanchez et al., 2022), the genetic etiology underlying this co-occurrence is not well understood (Akula et al., 2019; Cardozo et al., 2023). The categories “Eye development”, including “Visual system development”, and “camera-type eye development” were statistically significantly enriched in our dataset (q= 3.5E-02, Supplemental Table 3). Therefore, we re-examined the mutant lines with cleft palate using coronal optical sectioning of the micro-CT images in the IMPC database and found that the incidence of eye defects in our cleft palate mutants was about 25% (Table 1). The defects that we could phenotype were limited to morphological anomalies such as anophthalmia (the absence of one or both eyes), microphthalmia (small eyes), and obvious defects in the positioning of the optic cup (Figures 1-2, yellow font and an asterisk, Supplemental Table 1 for the list of specific embryos exhibiting both palatal and eye defects). According to the Centers for Disease Control and Prevention, the incidence of anophthalmia or microphthalmia in the human population is about 1 in 5100. The overall incidence of congenital ocular malformations varies between 0.36% and 4.7% depending on the study (Guarnera et al., 2024). Taking the upper probability of 4.7%, the binomial probability of 18 mutant lines displaying both the cleft palate and eye defects out of 72 examined is 3.86 x 10^-9^, indicating that the co-occurrence of cleft palate and eye defects in our dataset is not due to chance. In summary, our phenotyping effort has identified 18 genes regulating the development of the palate and the eyes, of which 8 (*Ankrd52, Cox7c, Pccb, Pfdn5, Podxl, Rgl1, Snx3,* and *Tmem115*) are novel regulators of palate development.

The NC is also important for ear development (Ritter and Martin, 2019), and the “Ear development” category was also among the statistically significantly enriched categories in our dataset (q=2.86E-02) (Supplemental Table 3). Among the genes in this category, *Maf* was a novel cleft palate gene. *Maf* encodes a group of transcription factors that can act as transcriptional activators or repressors. Of the four MAFs (*MAFA, MAFB, c-MAF, NRL*), *MAFB* is essential for the formation of inner ears, among its other functions, such as the development of pancreatic endocrine cells (Takahashi, 2021). Although not assigned to the “Ear development” category, *Runx1t1* is another novel cleft palate gene that may be involved in ear development. *Runx1t1* encodes a transcriptional corepressor implicated in the positive regulation of angiogenesis (Liao et al., 2017). In addition, Runx1t1 is commonly mutated in acute myeloid leukemia through its fusion with Runx1 (Grinev et al., 2021). Curiously, one case study reported a mutation in *Runx1t1* in a patient with a protruding ear (Huynh et al., 2012). In a recent study of birth defects associated with cleft palate with or without cleft lip, ear impairment was the third most common, after cardiac septation defects (Sanchez et al., 2022). The enrichment in the “Ear development” category in our dataset (Supplemental Table 3) suggests that screening for ear anomalies and hearing loss may be reasonable in patients with cleft palate.

The “Regulation of epithelial proliferation” category was also statistically a significantly enriched biological process (q= 2.04E-02, Supplemental Table 3). Among the 10 genes that fell into this category, 3 genes, *Cdk4, Celf1,* and *Klf7* were not previously known to regulate palate development. Our phenotyping analyses indicated that 1 of 6 homozygous *Celf1*-null mutants imaged by the IMPC exhibited a cleft palate (Figure 1). The binomial probability of cleft palate in this strain is p<0.004, which indicates that cleft palate did not occur simply by chance. *Celf1* (CUGBP Elav-like family member 1) is an RNA-binding protein regulating diverse aspects of mRNA biology, including mRNA alternative splicing, stability, and miRNA processing (Vlasova-St Louis et al., 2013).

*Celf1* knockout animals were previously generated on a mixed C57Bl6-129Sv background, and although ∼50% of the knockout animals did not survive past day 10 after birth, palatal development was not examined (Kress et al., 2007). Celf1 protein binds GU-rich sequences in introns as well as in 5’- and 3’-UTRs of mRNAs. The role of *Celf1* in palate development could be related to its ability to promote epithelial-to-mesenchymal transition (EMT) due to its ability to bind to and increase translation of mRNAs encoding EMT regulators such as *Snail1* (Chaudhury et al., 2016), which is important for palate development (Losa et al., 2018; Murray et al., 2007).

The enrichment in the “Respiratory system development” pathway was unexpected and especially intriguing as it contained the greatest number of novel cleft palate genes identified in our screen (Supplemental Table 3). 29 genes from our screen comprised this category, and 10 of them had not been associated with palate development before this work. These genes are *Acvr2b, Cdk4, Col27a1, Fgf15, Fndc3b, Maf, Pfdn5, Rgl1, Runx1t1,* and *Snx3. Col27a1* (a.k.a. type XXVII collagen) is expressed in the lung among other tissues (Pace et al., 2003) and is an evolutionarily conserved member of extracellular matrix proteins that belongs to a family of fibrillar collagens (Boot-Handford et al., 2003). The high conservation in vertebrates, the unusual domain composition relative to other fibrillar collagens, and the developmentally restricted expression of *Col27a1* suggested that this protein plays important developmental roles (Plumb et al., 2007). Mutations in the *Col27a1* gene have been associated with Steel syndrome, a rare autosomal recessive disease of the skeleton (Girisha et al., 2022; Gonzaga-Jauregui et al., 2020; Kotabagi et al., 2017). Among Steel syndrome patients, a pair of twin brothers was also diagnosed with cleft palate (Girisha et al., 2022). However, no other patients with *Col27a1* mutations or mouse mutants in *Col27a1* had palatal clefting (Plumb et al., 2011). Although only one of eight *Col27a1*-null IMPC mutants exhibited cleft palate (Figure 2), this incidence is statistically significant, p<0.005, excluding that cleft palate occurred purely by chance in this strain. However, the low incidence of cleft palate in *Col27a1*-null mutants may complicate further studies to understand the role of this protein in palatal shelf elevation or fusion. The enrichment in the “Respiratory system development” category suggests that cleft palate patients should be screened for potential defects in respiratory functions.

The enrichment in the “Kidney development” category (q= 8.22E-02) comprising *Acvr2b, Ap2b1, Cdkn1c, Ednra, Fgf10, Gdf11, Pygo2* genes was also surprising mainly because this set of genes did not include cilia genes, since defects in cilia development contribute both to craniofacial and kidney disorders (McConnachie et al., 2021; Moore, 2022). Published studies indicated that *Pygo2,* a component of canonical Wnt signaling is required for the development of both the palate and the kidney (Schwab et al., 2007), although the mechanism by which *Pygo2* mutants regulate palate development is still unknown. The enrichment in this category suggests the existence of common regulatory mechanisms mediating the development of the kidney and the palate in addition to those regulated by cilia.

In addition to biological processes, our data set was enriched for two pathways known to be involved in palatal development, “Regulation of activin receptor signaling pathway” (q= 3.51E-03) and “Regulation of smoothened signaling pathway” (q= 2.76E-02) were statistically significantly enriched. Interestingly, among the 13 genes in the “Regulation of activin receptor signaling pathway”, four were novel: *Acvr2b, Fndc3b, Arhgef18*, and *Fgf15*. Of these four genes, the homozygous-null *Arhgef18* mutants exhibited 100% penetrance of cleft palate (Table 1). *Arhgef18* gene encodes a RhoA guanine nucleotide exchange factor (GEF), also known as p114RhoGEF, that regulates RhoA activation and actomyosin contractility in multiple contexts (Beal et al., 2021; Nakajima and Tanoue, 2011; Tsuji et al., 2010; Xu et al., 2013). *Arhgef18*, thus, may be important for the fusion of the palatal shelves. As the two palatal shelves contact each other, the ectodermal epithelial cells, a.k.a. medial edge epithelial layers (MEE) that border the palatal mesenchyme, intercalate to form the midline epithelial seam (MES) (Lan et al., 2015). Activation of Rho and actomyosin contractility are essential for this intercalation and cell extrusion to form a continuous palatal tissue (Kim et al., 2015; Rosenblatt et al., 2001), and Arhgef18 could be an essential activator of RhoA in this process. Future conditional mutagenesis studies are necessary to test this hypothesis and to determine the precise tissues in which Arhgef18 functions during palatal development. This is especially important since recent studies found mutations in the *Arhgef18* gene in patients with sporadic cleft palate (El-Sibai et al., 2021).

Another homozygous null mutation, in the *Tctn3* (tectonic family member 3) gene, also resulted in penetrant cleft palate (Figure 1 and Table 1), albeit missed by the IMPC annotation pipeline. In humans, mutations in *Tctn*3 are associated with the Mohr- Majewski Syndrome, and a cleft palate was found in one case (Thomas et al., 2012). However, palatal development was not examined in *Tctn3*-null mutants generated before the IMPC screen (Wang et al., 2017); therefore, before our phenotyping screen, it was not clear whether *Tctn3* was required for the development of the palate. The *Tctn3* gene encodes a membrane protein that localizes to the primary cilium and is essential for the activation of the Shh signaling (Reiter and Skarnes, 2006; Wang et al., 2017). Ablation of *Tctn3* resulted in defective ciliogenesis (Wang et al., 2017).

Interestingly, ciliopathies (defects resulting in dysfunctional primary cilia) are common causes of craniofacial defects and underly several craniofacial syndromes, such as Joubert syndrome [OMIM 213300], Meckel-Gruber syndrome [OMIM 249000], Ellis-van Creveld syndrome [OMIM 225500], Oral–facial–digital syndromes types IX [OMIM 258865] and XIV (Adly et al., 2014; Thauvin-Robinet et al., 2014), and Weyers acrofacial dysostosis [OMIM 193530] (Moore, 2022; Reynolds et al., 2020). Consistent with this, our gene set enrichment analysis showed that the “Cilium assembly” category was detected in our dataset, although, not enriched at a statistically significant level (Supplemental Table 2). This category included the following genes: *Tctn3, Cep97, Inpp5e, Fuz*, and *Clasp1* (Supplemental Table 3), among which *Tctn3, Cep97*, and *Clasp1* are novel cleft palate genes identified in our screen (Figures 1-2). Cep97 localizes at the centrioles and regulates cilia assembly (Spektor et al., 2007). *Clasp1* encodes a plus-end microtubule-binding protein regulating the microtubule dynamics (Akhmanova et al., 2001; Maiato et al., 2005), an essential process for the biogenesis of cilia (Malicki and Johnson, 2017). Cilia are essential for the regulation of Shh and smoothened signaling (Goetz and Anderson, 2010), and our screen identified three additional genes (*Tctn3, Cep97,* and *Clasp1*) with joint roles in cilia, microtubule dynamics, and palatal development.

In summary, in addition to the identification of new genes involved in palatal development, our studies provide novel insights into the etiology of rare congenital defects such as those affecting both the eyes and the palate, and suggest that patients presenting with cleft palate may need to be followed up for anomalies in other organ systems, such as the heart, sensory nervous system, and the ear, as well as seemingly unexpected organs, such as the lungs and kidneys. Analyses of additional IMPC mutants in the future will provide important insights into the mechanisms of palatal development and identify additional genes, biological processes, and pathways important for palatal development and malformations. Our screen of IMPC mutants identified 45 novel genes regulating palatal development, providing fertile grounds for mechanistic studies in the future. Another value of such analyses lies in identifying unanticipated organs and pathways that could be affected alongside the palate in patients, allowing for a more thorough clinical follow-up and treatment of associated co- morbidities.

## Acknowledgments

These studies were supported by the National Heart, Lung, and Blood Institute of the National Institutes of Health (R01 HL103920, R01 HL134935, and R01 HL158049) to SA.

